# Modulating splicing in 5’ untranslated regions to treat rare haploinsufficient disease

**DOI:** 10.64898/2025.12.07.692584

**Authors:** Eloise S. Beer Wells, Laura De Conti, Hyung Chul Kim, Narjes Rohani, Kartik Chundru, Laura M. Watts, Ruebena Dawes, Yuyang Chen, Alexandra C. Martin-Geary, Michael J. Griffiths, Sam Scott, Rosemary A. Bamford, Jonathan Mill, Caroline F. Wright, Marco Baralle, Stephan J. Sanders, Nicola Whiffin

**Affiliations:** Big Data Institute, Nuffield Department of Medicine, University of Oxford, Oxford, OX3 7LF, UK; Centre for Human Genetics, University of Oxford, Oxford, OX3 7BN, UK; Institute of Developmental and Regenerative Medicine, Department of Paediatrics, University of Oxford, Oxford, OX3 7TY, UK; International Centre for Genetic Engineering and Biotechnology, Trieste, 34149, Italy; Clinical and Biomedical Sciences, University of Exeter Medical School, Exeter, EX1 2HZ, UK; Department of Psychiatry and Behavioral Sciences, UCSF Weill Institute for Neurosciences, University of California, San Francisco, CA 94158, USA

## Abstract

Rare genetic disorders collectively impact over 300 million people worldwide, yet around 95% have no specific treatments. For the many rare disorders caused by haploinsufficiency, effective therapies need to upregulate protein expression. However, therapeutic upregulation is often not straightforward. Increasing protein translation of the wildtype allele through inhibiting repressive upstream open reading frames (uORFs) has been proposed as a therapeutic approach for a few specific genes. The widespread success of steric-block antisense oligonucleotides (ASOs) for this purpose is, however, debated. Here, we explore an alternative approach, using splice-switching to exclude uORF-containing exons from the mRNA. Through a genome-wide computational screening approach, we identified 79 uORF-containing 5’UTR exons in haploinsufficient monogenic disease genes as candidate exon skipping targets. We demonstrate that removing the target exon significantly increased protein translation (between 1.4-5.5 fold) in a luciferase reporter assay for four of six prioritised target 5’UTR exons in neurodevelopmental disorder genes (*CTCF*, *GRIN2B*, *KRIT1*, and *TSC1*). Overall, this work supports the widespread application of 5’UTR exon skipping to boost translation of clinically relevant haploinsufficient genes.

## Introduction

It has been estimated that rare disorders collectively affect over 300 million people worldwide. They are associated with significant morbidity and mortality, but over 95% have no specific treatments^1^. Recent advances in oligonucleotide and CRISPR editing therapies are making personalised therapeutics that target specific genes or variants a reality for rare disease patients^2,3^. The ability of genome-targeted therapies to modify aspects of gene regulation underlies multiple successful therapies, including nusinersen for spinal muscular atrophy^4^, a therapy for Angelman’s syndrome which is showing promise in clinical trails^5^, and CASGEVY, a CRISPR therapy which induces foetal haemoglobin expression to alleviate sickle cell disease^6^. While it is relatively straightforward to down-regulate genes therapeutically, for example with gapmer antisense oligonucleotides (ASOs), upregulating protein levels in diseases of haploinsufficiency is considerably more difficult^7^, often requiring a bespoke approach for every gene.

Upstream open reading frames (uORFs) are negative regulators of translation that are found within five prime untranslated regions (5’UTRs)^8^. They are highly conserved and are predicted to occur in half of human protein-coding genes^9^. The presence of a uORF disrupts ribosome scanning through the 5’UTR, limiting ribosome progression to translating the downstream canonical protein sequence. uORFs have been shown to reduce translation efficiency of the downstream coding sequence (CDS) by 30-80%^8^. The translation of uORFs can be detected by ribosome profiling (ribo-seq), a technique that captures and sequences ribosome-bound RNA fragments^10^.

Removing uORF-mediated translational repression has been proposed as an approach to upregulate protein levels therapeutically. Early studies showed that this could be achieved using steric-blocking ASOs directly targeted to uORFs^11^, with recent success in upregulating *JAG1* in a mouse model^12^. Others have, however, failed to replicate the approach^13^. It is conceivable that a steric-block ASO bound to a target 5’UTR may itself block ribosome scanning and result in reduced rather than increased downstream protein translation. A further study used ASOs to disrupt an RNA secondary structure which promotes translation from an uORF, thus upregulating the canonical protein^14^.

Splice switching ASOs (SSOs) can modify exon inclusion in mRNA transcripts. They have been developed and approved to treat Duchenne Muscular Dystrophy (DMD) and Spinal Muscular Atrophy (SMA)^15^. Over one third (37.7%) of human protein-coding genes have multiple 5’UTR exons^9^. In these genes, SSOs could be used to modify 5’UTR sequence and structure and hence translational regulation. For example, inclusion of a cryptic 5’UTR exon in *PRNP* which contains a uORF can reduce prion protein (PrP) expression^16^.

Here, we propose that for a subset of genes, SSOs could promote skipping of a uORF-containing 5’UTR exon to upregulate translation from the functional wildtype allele of the gene in diseases of haploinsufficiency. The success of this approach was recently demonstrated for a single target gene, *BDNF*^17^. We describe a systematic approach to identify genes where uORF-containing 5’UTR exon skipping is tractable and release a resource of candidate target exons. We functionally validate six prioritised target exons, demonstrating a therapeutically relevant increase in downstream protein translation when excluding uORF-containing exons in four of these. Together, these results highlight the promise of 5’UTR exon skipping as a therapy for a subset of haploinsufficient disorders.

## Results

### Many genes have skippable 5’UTR exons containing an upstream start codon

We developed a genome-wide approach to identify candidate skippable 5’UTR uORF-containing exons (**Figure 1**). We define ‘skippable’ exons as those which encode the 5’UTR and are flanked by introns. Hence, the first 5’UTR exon, and any exon which contains canonical coding sequence, are not considered ‘skippable’. Using the Matched Annotation from NCBI EMBL-EBI (MANE) v1.0 transcript dataset we identified 2,210 ‘skippable’ 5’UTR exons in 1,572 genes.

**Figure 1:**
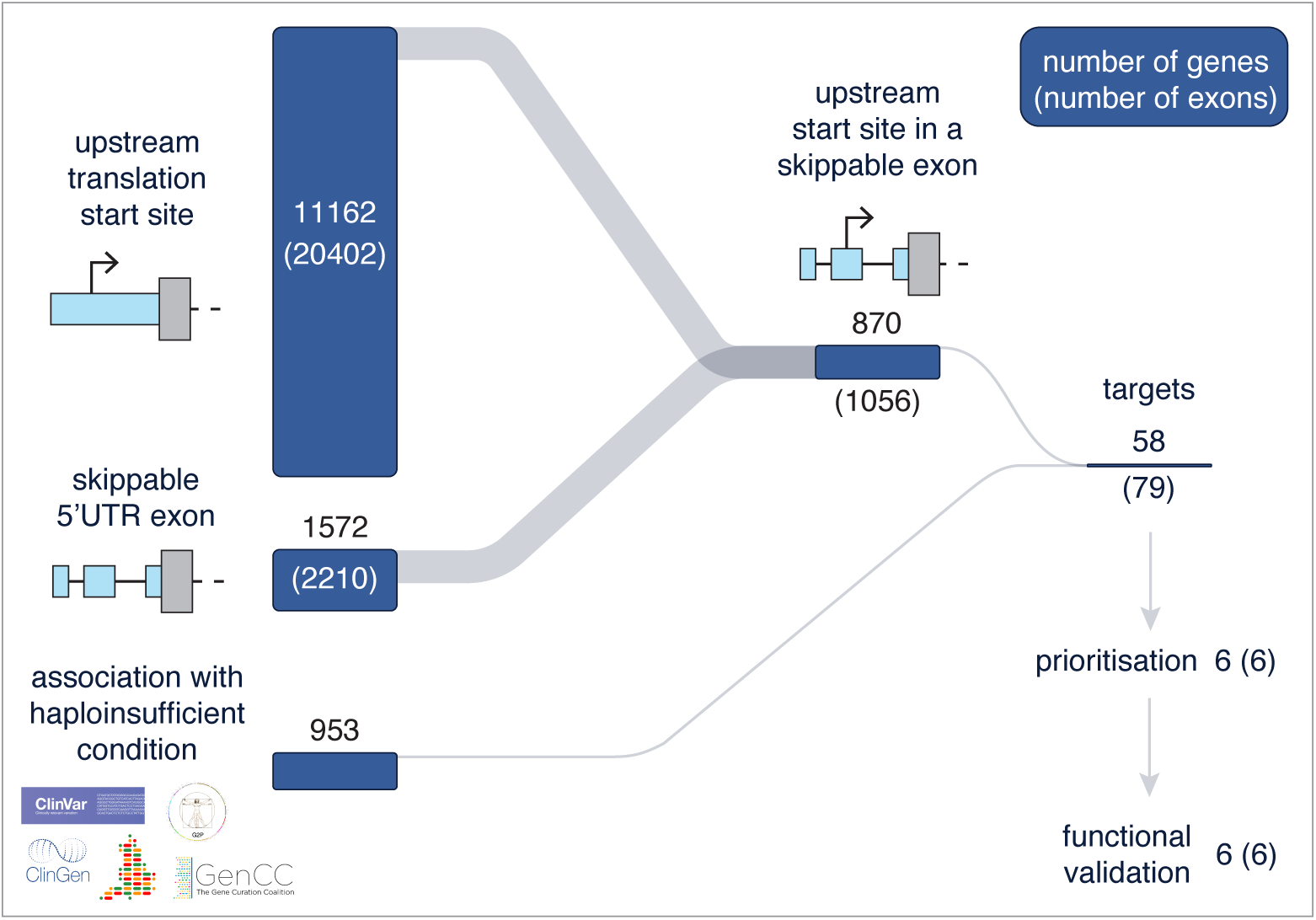
Overview of the target identification process. For each step, the number of genes in each category is shown in or above the bar, the number of exons in each category is displayed in brackets. Upstream translation start sites were identified from nine ribo-seq studies. Skippable 5’UTR exons were defined as those flanked by introns in MANE v1.0, which could feasibly be excluded by disrupting the splice site. These exons were cross-referenced against haploinsufficient genes collated in clinically-curated gene-disease databases. The genes were then prioritised based on estimated prevalence of the associated disorder (calculated based on ClinVar^18,19^, Kaplanis et al.^20^, and diagnoses from the National Genomics Research Library), the 5’UTR structure, and the strength of disease association (see **Methods**). Based on this prioritisation, six target genes/exons were chosen for functional validation.

We collated 60,786 upstream start codons, in 11,162 genes, identified from nine ribo-seq studies performed across 13 human tissues, primary cells and cell lines, including brain, heart and skeletal muscle (see **Methods, Supplementary Figure 1**). Of these 60,786 upstream start codons, 2,187 (3.6%) were within 1,056 skippable 5’UTR exons, in the transcripts of 870 unique genes (**Figure 1**).

We filtered the candidate exons to those within genes that are a known cause of haploinsufficient disorders (i.e., with a monoallelic loss-of-function mechanism) defined using GenCC^21^, PanelApp^22,23^, G2P^24,25^ and ClinVar^18,19^ (see **Methods**). This yielded 79 candidate exons in 58 genes (**Figure 1; Supplementary Table 1**). We prioritised six target exons in six different genes for further characterisation and functional validation (**Table 1**). Prioritisation was based on the strength of association with a haploinsufficient developmental disorder, the estimated prevalence of the disorder, and the overall 5’UTR structure and predicted impact of exon skipping on translation (see **Methods**).

**Table 1:** Details of the six target exons selected for experimental investigation. The consequence of 5’UTR exon exclusion is illustrated in **Supplementary Figure 3**. All exons are defined by MANE select transcripts. All targets exclude at least one high confidence uORF; defined as those that initiate at an ATG start codon and/or are identified by more than two ribo-seq datasets. Additional information for all candidate exons is in **Supplementary Table 1**. *variants impacting either the splice acceptor or splice donor site of the target exon **defined as a uORF which has a strong likelihood of prohibiting translation reinitiation at the CDS: either a uORF which creates an intercistronic region of less than 50 ntds to the CDS, or an overlapping uORF (uoORF). 100kGP: the Genomics England 100,000 genomes project from the National Genomics Research Library, UKBB: UK BioBank, GenCC: the gene curation coalition, G2P: Gene2Phenotype, LOEUF: loss-of-function observed/expected upper bound fraction.

| target information |  |  |  |  | consequence of exon exclusion |  |  |  | disease association |  |  | estimated prevalence |  |  |
| --- | --- | --- | --- | --- | --- | --- | --- | --- | --- | --- | --- | --- | --- | --- |
| gene symbol | total 5'UTR exons | 5'UTR length (ntds) | target exon number | target exon length (ntds) | UKBB variants at exon splice sites* | Change in no. high confidence uORFs | no. high confidence uORFs remaining | Removes a strongly repressive high confidence uORF** | GenCC/G2P definitively associated conditions | G2P variant consequence | LOEUF | ClinVar pathogenic variants | Kaplanis<sup>20</sup> pLoF variants | 100kGP diagnoses |
| *ANKRD11* | 3 | 461 | 2 | 83 | 1 | -1 | 2 | yes | KBG syndrome | decreased gene product | 0.13 | 561 | 111 | 62 |
| *CTCF* | 3 | 325 | 2 | 115 | 4 | -1 | 0 | yes | Intellectual disability | absent gene product | 0.20 | 45 | 9 | 7 |
| *GRIN2B* | 3 | 708 | 2 | 427 | 1 | -3 | 1 | yes | Epileptic encephalopathy, Intellectual developmental disorder | altered gene product structure, absent gene product | 0.10 | 103 | 11 | 32 |
| *KRIT1* | 4 | 545 | 2 | 268 | 0 | -5 | 1 | no | Cerebral cavernous malformation | absent gene product | 0.69 | 310 | 0 | 4 |
| *TBL1XR1* | 3 | 483 | 2 | 74 | 0 | -3 | 0 | yes | Intellectual disability with autism spectrum disorder/complex neurodevelopmental disorder, Pierpont syndrome | absent gene product, altered gene product structure | 0.17 | 33 | 7 | 3 |
| *TSC1* | 3 | 217 | 2 | 61 | 1 | -1 | 2 | no | Tuberous sclerosis | absent gene product/altered gene product structure | 0.23 | 653 | 2 | 8 |

### Prioritised target exons are included in brain expressed transcripts through development

For the exclusion of a uORF-containing exon to be an effective treatment for a rare disorder, the exon must be present in a high proportion of transcripts in phenotypically relevant tissues and at appropriate developmental stages. For each of our experimentally validated genes we investigated splicing patterns in the brain using both short- and long-read RNA-sequencing data.

Using short-read, bulk tissue, Ribo-Zero RNA-seq data from 116 prenatal and 60 postnatal prefrontal cortex samples from BrainVar^26^, we compared the expression of the target 5’UTR exon with a length-matched 5’ portion of the coding sequence (CDS). For four genes (*ANKRD11, TBL1XR1, TSC1, KRIT1*), the relative expression of the target exon and CDS reference portion remains constant throughout development (**Figure 2A, Supplementary Table 2**). For all genes, CDS exon expression was higher than that of the 5’UTR, which could be reflective of either alternative splicing or 3’ bias in the dataset. The highest CDS exon expression relative to the 5’UTR target exon was seen for *CTCF* at 3.80-fold and the lowest was *TBL1XR1* at 1.32-fold.

**Figure 2:**
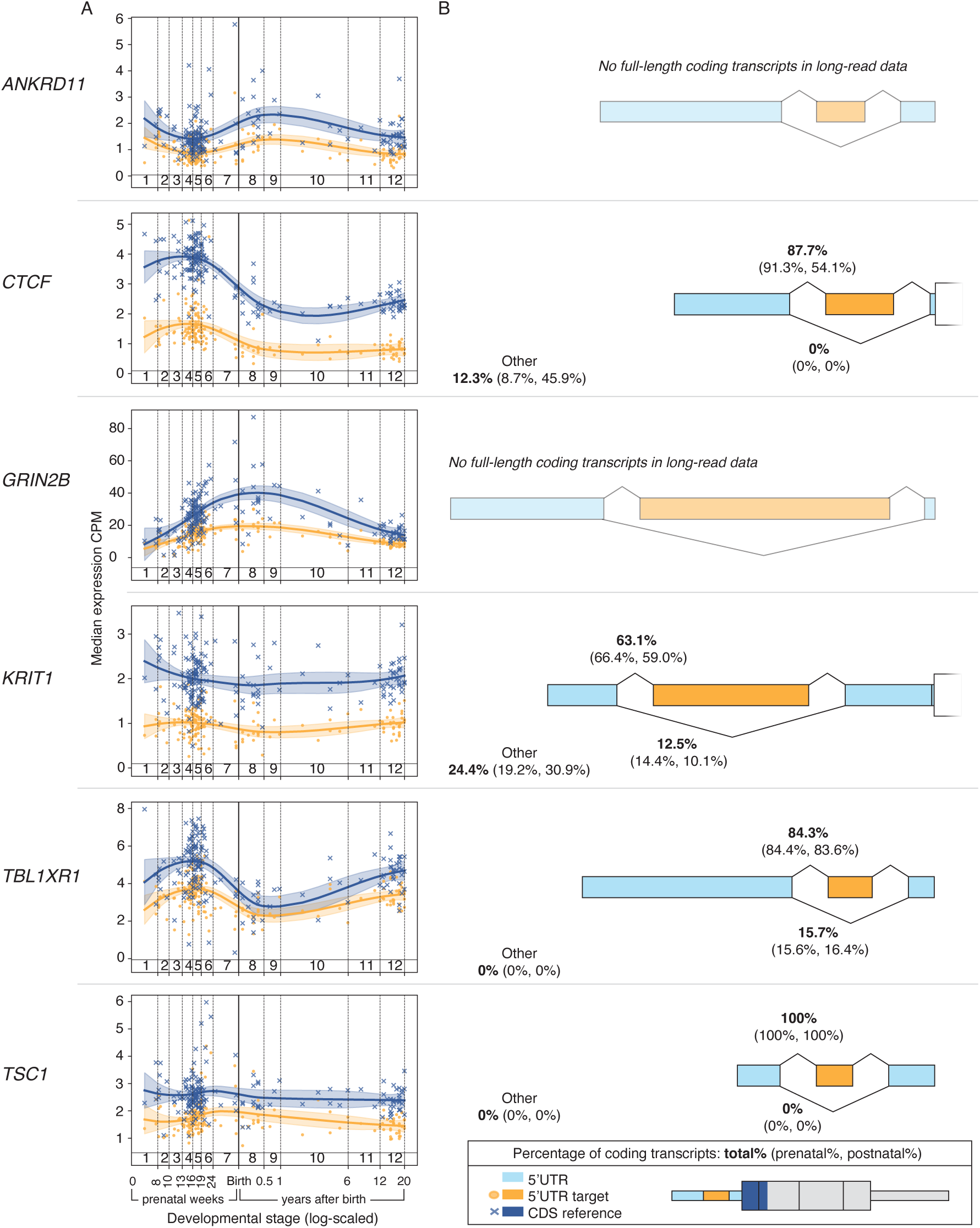
Alternative splicing of six prioritised target exons. **A)** Exon expression across development in short-read RNA-sequencing data from human prefrontal cortex in BrainVar^26^. Median log-transformed expression across samples is displayed for the first canonical coding exon (dark blue) and the 5′ UTR target exon (orange). An estimated scatterplot smoothing (LOESS) curve was fitted to the data to capture expression trends over time, with shaded areas indicating the 95% confidence interval**. B)** Characterisation of alternative splicing in long-read RNA-sequencing data in human prefrontal cortex^27^. Schematics for each gene show the proportion of full-length coding transcripts (see **Methods** and **Supplementary Figure 2**) that contain or skip the target 5’UTR exon. For *ANKRD11* and *GRIN2B,* no full-length transcripts were detected. These transcripts are the longest of the six target genes (9,301 nucleotides and 30,609 ntds respectively). For all other genes, the target exon was present in the majority of transcripts (>63%).

We further characterised full-length transcript expression using long-read RNA-sequencing data from 26 prenatal and 21 postnatal frontal cortex samples^27^. To account for 3’ bias in these data, we restricted our analysis to transcripts with a transcription start site overlapping a CAGE peak from the FANTOM5 resource and that included at least 80% of the MANE CDS. Four of our six prioritised genes had one or more transcript fitting this definition (**Supplementary Figure 2**). Conversely, no such transcripts were detected for the two largest genes, *GRIN2B* and *ANKRD11*. For the four genes with full-length transcripts, we calculated the proportion of transcripts that: (1) used the full-length 5’UTR including the target exon, (2) excluded the target exon but maintained the upstream and downstream splice-sites, or (3) had a different 5’UTR structure. The lowest target exon inclusion in expressed transcripts was observed for *KRIT1*, at 63.1%, rising to 100% for *TSC1*. The proportion of reads from transcripts where the target exon is skippable did not change substantially across development, except for *CTCF*, where inclusion of the target exon was significantly higher in prenatal compared to postnatal samples (91.3% vs 54.1%, Fisher’s exact test, odds ratio = 0.11, p = 5.31x10^-8^, **Figure 2B, Supplementary Figure 2**).

### Investigating additional features that influence translational regulation

To gain a more comprehensive picture of 5’UTR mediated regulation in each of our six selected genes, we predicted the occurrence of stem loops and G-quadruplex structures, and annotated RNA binding protein (RBP) binding sites, within the full-length 5’UTR of each gene. We used RNAfold^28^ to identify 116 predicted stem loops within the 5’UTRs of the six genes (median = 19.5, range 10–31). Exclusion of target exons had a minor impact on the number of stem loops, with no change in the predicted number for three targets (*GRIN2B, TBL1XR1, TSC1*). For stem loops that were not removed (n = 81) half (47%) changed in length (-5ntds to +38 nucleotides). We identified 20 predicted non-overlapping G-quadruplexes in the 5’UTRs of the genes (median=3, range 2–6), three of which would be disrupted by the exclusion of the target exon (in *GRIN2B*, *CTCF*, and *TSC1*; **Supplementary Figure 3**)^29^. We identified 19 RBP binding sites within the 5’UTRs of the genes (median = 2, range 0-13)^30^, two of which (in *ANKRD11* and *CTCF*) overlap a target exon and would be removed if it were skipped (**Supplementary Figure 3**). There were no experimentally confirmed microRNA (miRNA) binding sites or internal ribosome entry sites (IRES) within any of our target gene 5’UTRs^31,32^. In summary, we predict that exclusion of each of the target exons would have minimal impact on other 5’UTR regulatory features.

### Exclusion of target exons significantly increases translation

To test whether the exclusion of the prioritised target exons resulted in increased translation, we designed a dual-glo luciferase assay. For each target exon, we cloned three 5’UTR constructs upstream of the *Renilla* luciferase reporter: 1) full-length wildtype 5’UTR, 2) the 5’UTR without the target exon, 3) full-length 5’UTR with all ATG codons in the target exon changed to ATC (**Figure 3A, Supplementary Table 3**). The latter construct was included to elucidate whether any detected reduction in translation was likely attributable to removal of the uORF(s) versus disruption of the other regulatory features described above.

**Figure 3:**
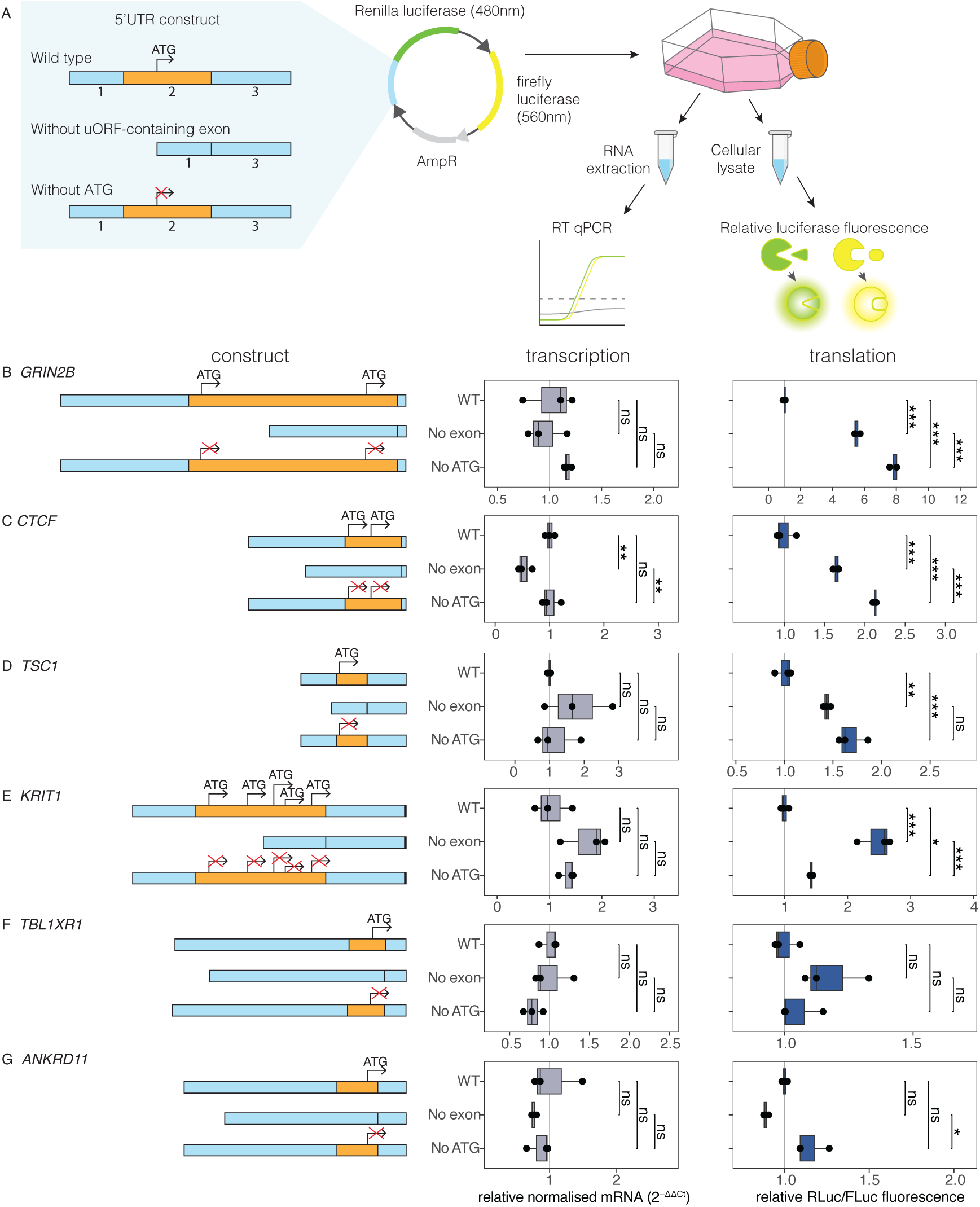
Exclusion of uORF-containing 5’UTR exons can boost translation. **A)** Schematic illustration of the experimental design. For each gene, three constructs were tested in a dual luciferase reporter system: (1) full-length wildtype 5’UTR, (2) 5’UTR without the candidate exon, (3) full-length 5’UTR with all ATGs in the target exon changed to ATC. Relative renilla luciferase to firefly luciferase was used to assess changes in translation and RT-qPCR was used to detect changes in mRNA levels. **B-G**) Construct illustrations, RT-qPCR, and luciferase assay results for **B)** *GRIN2B*, **C)** *CTCF*, **D)** *TSC1*, **E)** *KRIT1*, **F**) *TBL1XR1*, and **G)** *ANKRD11*. Data were analysed using one-way ANOVA, followed by Tukey’s HSD multiple comparison test. Asterisks denote statistical significance ***<0.001, **<0.01, *<0.05, ns = non-significant.

For four of the six genes (*CTCF, GRIN2B, KRIT1,* and *TSC1*), luciferase activity was significantly increased following removal of the target exon, with observed changes of 1.6- to 5.5-fold. For each of these genes, luciferase activity was also significantly increased in constructs with full-length 5’UTR but with ATGs in the target exon changed to ATC with observed changes of 1.4- to 7.9-fold (**Figure 3B-E**). For *TSC1* there was no significant difference between the effect of removing exon 2 to that of mutating ATG sites (Tukey HSD test, mean difference = 0.034, p adj = 0.068). For *CTCF* and *GRIN2B* the effect of removing the ATGs was significantly greater than removing the target exon (Tukey HSD test, *CTCF*: mean difference = 0.24, p adj = 7.72x10^-4^, *GRIN2B*: mean difference = 0.31, p adj = 1.13x10^-5^). Conversely, in *KRIT1*, removing the target exon had a significantly greater effect than removing the ATGs (Tukey HSD test, mean difference = 0.053, p adj = 5.97x10^-4^). We observed no significant change in luciferase activity following the removal of the target exon, or mutation of ATGs within that exon, for either *ANKRD11* or *TBL1XR1* (**Figure 3F-G**).

### Protein translation can be reduced by uORFs in an additive manner

For five of the six prioritised gene targets, there are multiple high confidence uORFs within the 5’UTR. These uORFs may act redundantly and hinder the upregulation of gene expression through uORF skipping^17^. To examine this further, we picked a single target gene, *TSC1*. Haploinsufficiency of *TSC1* causes tuberous sclerosis, a neurocutaneous condition characterised by multisystem benign tumour growth, including brain lesions, alongside epilepsy, cognitive impairment, developmental delay and autism spectrum disorder^33^. The 5’UTR of *TSC1* contains three exons, each containing a single ATG start codon, each of which encodes a uORF that is between 21-63 bp long and two of which are detected in multiple ribo-seq datasets (**Supplementary Figure 3**). In our above assay, removal of the central 5’UTR exon resulted in 1.4-fold increase in translation, with removal of the ATG in this exon leading to 1.7-fold increase.

To investigate uORF-mediated translational regulation in *TSC1*, we systematically mutated each ATG to ATC individually, and in combination, within our dual-glo luciferase assay system. We observed an additive effect of mutating the uORF start codons: a 1.3-1.9-fold increase in luciferase activity when mutating each single ATG, 2.2-3.0-fold upregulation when mutating two stop codons, and the greatest upregulation of 5.4-fold was observed when all three uORF start codons were mutated (**Figure 4**).

**Figure 4:**
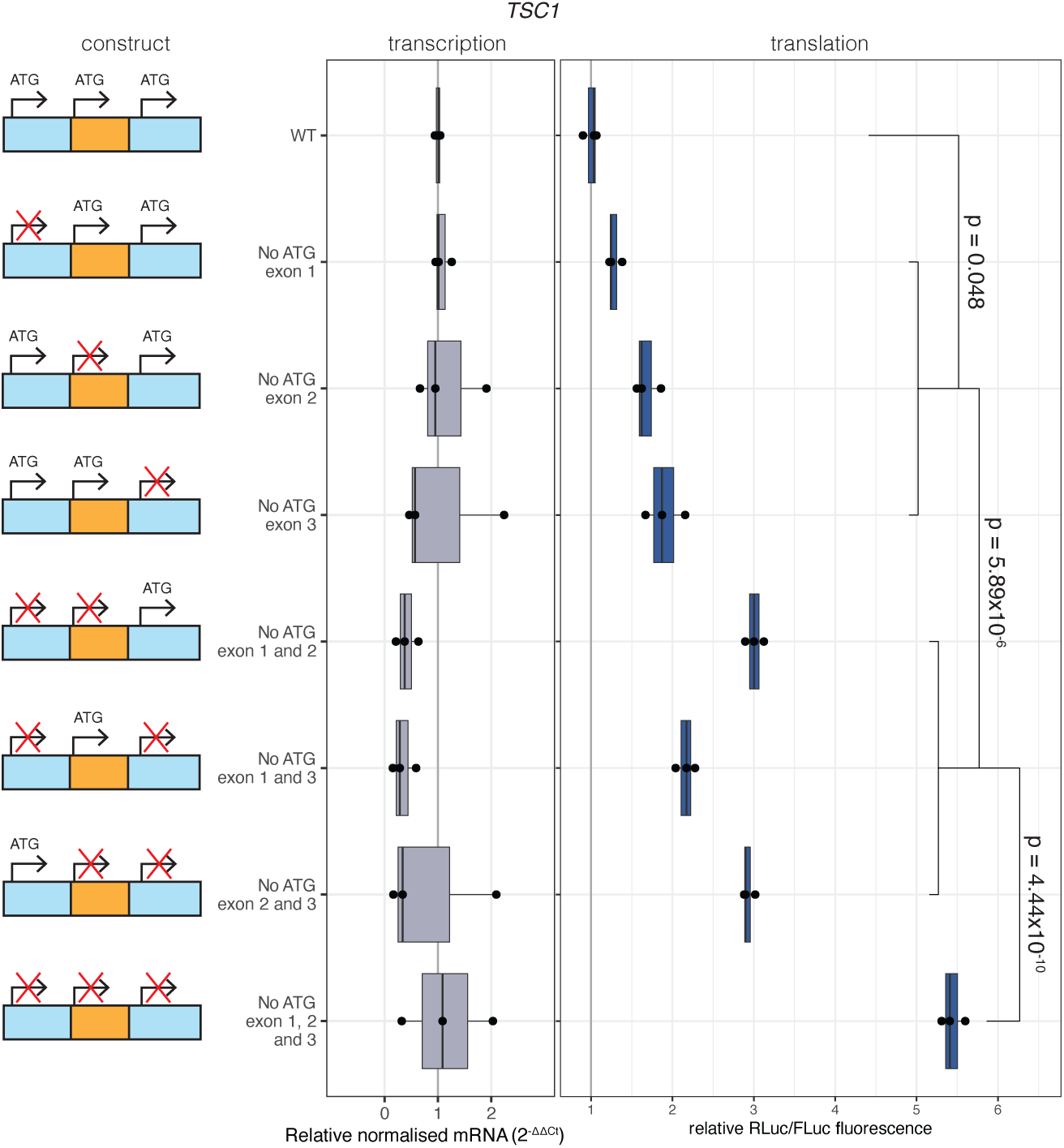
uORFs have an additive effect on translation repression in *TSC1*. Construct illustrations, RT-qPCR, and luciferase results for a systematic test of the effect of uORFs on luciferase translation. There was a significant effect of construct on normalised reporter fluorescence. (ANOVA: F=320.66; d.f.= 8,18; p= 2.20x10^-16^). Results of statistical analysis for all comparisons are in **Supplementary table 6**. There was no significant effect of construct on quantity of mRNA (ANOVA: F=.3169; d.f.= 8,21; p=0.289).

## Discussion

Here, we developed a computational approach to determine which haploinsufficient disorders are potentially treatable with a 5’UTR exon skipping approach to upregulate protein translation. We describe a framework to identify target exons, release an open-source resource of these tractable exons, and show the utility of this approach through further dissection and functional validation of six prioritised targets.

Using luciferase reporter constructs, we demonstrated that exclusion of uORF-containing 5’UTR exons resulted in 1.4-to-5.5-fold upregulation in translation of four of six target genes. As haploinsufficiency is a result of loss-of-function of a single allele, a ∼2-fold increase in protein translation could reasonably be expected to compensate. However, for some genes, even a modest increase in translation could be therapeutically beneficial. For example, only a partial correction in *SCN1A* protein production rescues the epileptic phenotype in haploinsufficient mouse models of Dravet syndrome^34^. Conversely, for some genes both loss- and gain-of-function lead to severe phenotypes. For these genes the level of upregulation driven by a therapy might need to be tightly controlled. An example is *GRIN2B*, where we observed 5.5-fold upregulation when removing the target 5’UTR exon, the largest of any of our prioritised targets. For such genes it will also be critical to be certain of the mechanism underlying the disease-causing variant in each patient prior to treatment.

In addition, we determined the extent to which any change in translation was due to the uORF, versus other features within the exon. We mutated all ATG codons within the target exon to ATC (a codon not known to drive appreciable levels of translation). In all four genes with significant increases in translation in response to exon skipping, there was also significant increases in translation in response to mutation of ATGs within that exon. However, in only one case, *TSC1*, there was no significant difference between these two approaches, pointing to the complexity of translational regulation, with multiple features working in tandem to control protein expression. Interestingly, for two of the four genes in which we observed significant increases in translation (*GRIN2B, CTCF*), mutation of the uORF start codons led to a significantly greater increase in protein translation than exclusion of the exon they reside within. Therefore, targeting the uORF start codons with single base editing techniques could also be an effective therapeutic approach.

To further investigate the complexities of uORF-mediated translational regulation, we made single and combinatorial edits to uORF start codons in a single gene target, *TSC1*. Our results show that uORFs in this gene act additively and, when a single uORF is excluded, those more proximal to the CDS have a greater impact on translation, in accordance with previous findings^35^. This highlights how the impact of removing a uORF, or the exon it resides within, is dependent on the context of the whole 5’UTR.

In response to target exon skipping, a uORF upstream of the target exon may have a greater inhibitory impact on translation through increased proximity to the CDS, greater distance to the next inframe stop codon, or the loss of a stop codon in the 5’UTR. This could also explain why exclusion of two of the exon targets, in *ANKRD11* and *TBL1XR1*, did not result in increased translation: in each case, a uORF further upstream would be brought closer to the CDS, potentially maintaining translational repression. In addition, a uORF downstream of the target exon could act as a regulatory bottleneck. In both *ANKRD11* and *TBL1XR1,* additional ATGs exist downstream of the target exon that could be translated at higher levels if they become more accessible to scanning ribosomes. Indeed, when we mutated the ATG codons in exon 3 of *ANKRD11*, in addition to those in exon 2, we observed a 2.8-fold increase in translation compared to WT (**Supplementary Figure 4**). While we did analyse the 5’UTR structure during target prioritisation, the impact of these changes on downstream translation remain contextually unique and therefore difficult to predict^36^. These predictions may be improved in the future, alternatively, large-scale functional assays that simultaneously test the removal of all skippable 5’UTR exons on downstream translation would be incredibly valuable for target identification.

Our study has multiple limitations. While expansive, the catalogue of uORFs used in this study is not comprehensive, owing to the incomplete sampling of human cell types and developmental states. We therefore may miss some translated uORFs and have an incomplete view of the uORF makeup across each entire 5’UTR. Conversely, ribosome profiling datasets are currently noisy, with many identified uORFs potentially not translated at high levels. ORF calling pipelines might also struggle to accurately identify the correct uORF start codon. Hence, functional validation of each target uORF and its effect on downstream CDS translation is important. To combat these limitations, we restricted target selection to uORFs identified across multiple datasets, hypothesising that these are more likely to represent true translation events. Here, our functional validation used a reporter construct within a model cell line that simulates target exon exclusion in 100% of transcripts. Therapeutic application will depend upon both the efficacy of exon exclusion by SSOs or gene editing technologies, and comparable upregulation of the canonical protein product being observed in phenotypically-relevant cell types and endogenous contexts. Further, our analysis of RNA-sequencing datasets indicated incomplete inclusion of target exons in brain-expressed transcripts. Hence, the level of translational upregulation achieved *in vivo* will likely be lower than what we detect in our luciferase assay. Future experimental validation of target exons should be performed on native 5’UTR sequences in a disease relevant cell-line. Finally, the interaction of structural elements of the 5’UTRs with uORFs influence translational regulation^14,37^. Other elements such as RBPs are also known to influence translation. Our ability to predict the occurrence of these elements, and their impact on CDS translation is currently limited.

Overall, we present a systematic strategy to identify haploinsufficient human disease genes which may be upregulated via an exon skipping strategy through the exclusion of uORFs. This could be achieved by using SSOs, or genome editing to disrupt splice-sites. SSOs have the benefit of reversible and dosage sensitive effects, helping to derisk future irreversible genome-editing approaches. Importantly, SSOs are a variant-agnostic approach to treating haploinsufficiency; this therapeutic mechanism should be accessible to all individuals with haploinsufficiency caused by any variant resulting in loss-of-function in a gene with a uORF-containing skippable exon. This work adds to the growing number of approaches to achieve therapeutic upregulation, underscoring the promise of genome targeted therapies for the treatment of previously intractable rare disorders.

## Methods

### Data and code availability

A snakemake workflow for uORF start codon collation, haploinsufficient gene identification, target identification and prioritisation is available here: https://github.com/Computational-Rare-Disease-Genomics-WHG/skip_uORFs/

RNA structure was predicted using RNAfold^28^. Code available here: https://github.com/Computational-Rare-Disease-Genomics-WHG/RNA_structure

### National Genomics Research Library data

Research on the de-identified patient data used in this publication can be carried out in the Genomics England Research Environment subject to a collaborative agreement that adheres to patient led governance. All interested readers will be able to access the data in the same manner that the authors accessed the data. For more information about accessing the data, interested readers may contact or access the relevant information on the Genomics England website: https://www.genomicsengland.co.uk/research.

### Identification of transcripts with skippable exons

Transcripts were defined using MANE v1.0 (available at https://ftp.ncbi.nlm.nih.gov/refseq/MANE/MANE_human/release_1.0/). Skippable exons were defined as any exon that is flanked by introns in the 5’UTR. This excludes the first 5’UTR exon, and typically the final exon of the 5’UTR, unless there is an intron between the end of the 5’UTR and the start of the CDS.

### Identification of upstream start codons using ribo-seq

We collated ORF calls from nine published ribo-seq studies (**Supplementary Table 5**). Where necessary, we used liftover to transform genome coordinates to GRCh38 and adjusted coordinates such that canonical ORFs aligned with their annotations in UCSC genome browser (1-based). The sequence of the start codon was retrieved against the hg38 reference. Where sources reported the sequence of the start codon, ORFs with a start codon sequence mismatch were excluded.

### Identification of genes associated with autosomal dominant haploinsufficiency

We used ClinVar^18,19^, GenCC^21^, Gene2Phenotype (G2P)^24,25^, ClinGen gene-disease validity^38^, and Genomics England^22,23^ to identify 953 genes associated with dominant loss-of-function haploinsufficient disorders. The list includes (1) all ‘Definitive’ and ‘Strong’ genes which are tagged as ‘monoallelic’ and ‘absent gene product’ in at least one G2P dataset (cancer, cardiac, developmental, eye, skeletal, and skin^39^, (2) genes with haploinsufficiency scores of 3 ‘Sufficient Evidence’ from the ClinGen dosage sensitivity^40^, (3) genes that were flagged as ‘Green’ and ‘MONOALLELIC’ in PanelApp^41^, and (4) genes with ‘Definitive’ or ‘Strong’ GenCC classifications and with a mode-of-inheritance of ‘dominant’^42^. For PanelApp and GenCC, where there was no information on variant mechanism available, genes were filtered to those that also had at least eight pathogenic/likely pathogenic predicted loss-of-function variants in ClinVar^43^.

### Prioritising genes and exons for follow-up

Six exon targets were selected for functional validation based on their strength of association with developmental disorder, estimated prevalence, and predicted consequence of exon skipping on translation. Specifically, they met these criteria: 1) Exon exclusion does not cause the extension of other high confidence uORFs such that they stop near to (<50 ntds), or overlap with, the CDS; 2) The gene is definitively associated with an autosomal dominant developmental condition by G2P or GenCC; 3) The gene is in the top 60% of counts of ClinVar pathogenic variants, Kaplanis pLoF variants or National Genomics Research Library’s 100,000 genomes project data (100kGP) diagnoses; 4) Exon exclusion results in a removal of a high confidence uORF which stops near the CDS (<50ntds), removal of a high confidence ouORF, or removal of three or more high confidence uORFs. In addition, *TSC1* was selected due to its unique 5’UTR structure for interrogating the impact of multiple uORF perturbations.

### Strength of association with developmental disorder

We prioritised genes with a strong or definitive association with a developmental disorder, with autosomal dominant inheritance. We identified such genes within the G2P DD panel^39^, and by cross-referencing GenCC associations with MONDO hierarchies (0021147 disorder of development or morphogenesis, 0700092 neurodevelopmental disorder or 0005503 developmental disorder of mental health,^42,44^.

### Estimate of prevalence

We used 100kGP v18, and loss-of-function variant counts from ClinVar (see above), and in Supplementary Table 1 from Kaplanis *et al*^20^ to estimate prevalence. For 100kGP, diagnostic data was obtained from the Genomics England ‘exit questionnaire’ (12,024 unique variants in 9,841 individuals) and ‘diagnostic discovery table’ (2,394 unique variants in 2,452 individuals respectively) via the RLabkey API. Where required, variants were remapped to GRCh38 using the UCSC LiftOver tool^45^. Variants were filtered to include only those with an annotation of “pathogenic”, “likely pathogenic”, or that had been identified in the “diagnostic discovery” table (8,145 unique variants in 8,709 individuals), and that mapped to a single gene (5,752 unique variants in 6,095 individuals). This resulted in a set of 5,752 pathogenic/likely pathogenic variants in 6,056 unique individuals

### Predicted consequence of exon skipping on the 5’UTR and translation

Exon exclusion can cause an extension of upstream uORFs due to stop-loss, for instance the third exon of *PAX6*^46^. For this reason, sixteen exon targets predicted to cause uORF extensions were removed from consideration. Of the remainder, exon targets were only considered if a high confidence uORF initiated within them (n = 52). uORFs which are closer to the CDS, or overlap with the CDS, are highly repressive, so we prioritised exon targets which would exclude a uORF within 50bp of the CDS, or an overlapping upstream ORF. In addition, hypothesising that removing multiple uORFs would have a greater impact on translation, we prioritised exon targets which would exclude >3 high confidence uORF start codons. 3) We also included exon targets with at least one probable splice-disrupting variant in UK Biobank (n=44), hypothesising that this indicates that skipping the target exon is tolerated. Data were accessed under UK Biobank application number 81050. UK Biobank protocols were approved by the National Research Ethics Service Committee.

### Identification of RNA stem loops, G-quadruplexes, IRES, RBPs & miRNA binding sites

Stem loops were identified using RNAfold^28^. A 60ntd sliding window was applied to the 5’UTR of each gene, plus the first 50ntds of the CDS, as determined using 5’UTR annotations from MANE and human reference genome hg38 (available at http://hgdownload.soe.ucsc.edu/goldenpath/hg38/bigZips/latest). Stem loops which were located wholly within another stem loop were excluded. G-quadruplexes were identified with QGRS mapper^29^. RBPs were identified with RBP-Tar^30^ available at https://rbp-tar.biodata.ceitec.cz/. IRES were searched with IRES base^32^. The binding of microRNA to the 5’UTR was investigated with miRStart 2.0^31^, but no miRNA binding sites were observed.

### Analysis of human cortex long-read RNA-seq data

Oxford Nanopore Technologies long read RNA sequence data was generated from 26 prenatal and 21 postnatal frontal cortex samples are described in Bamford *et al.* 2024^27^. The transcripts were annotated using SQANTI3^47^ and CPAT was used to predict the open reading frame and coding potential of the transcripts^48^. Transcript isoforms were considered protein-coding if the CPAT coding probability was the recommended >0.364, the open reading frame had a canonical stop site, 80% of the CDS as defined by MANE was included, and the start of the transcript was within 500bp of a CAGE peak^49^. From these reads, we calculated the proportion in which the target exon was flanked by the first and third 5’UTR exon and deemed skippable (**Figure 2, Supplementary Figure 2**).

### Analysis of exon expression in the human cortex with short read RNA-seq data

Short read RNA sequence data from 116 prenatal and 60 postnatal dorsolateral prefrontal cortex samples are described in Werling *et al*^26^. Exon-level expression was quantified from bulk RNA-seq data using exon annotations derived from DEXSeq. For each sample, read fragments overlapping the 5′ UTR target exon region, and a length-matched portion at the 5’ end of the CDS, were counted and median of those fragments were calculated. Expression values were normalized to counts per million (CPM) to account for differences in sequencing depth. An estimated scatterplot smoothing (LOESS) curve was fitted to the data to capture expression trends over time, with shaded areas indicating the 95% confidence interval of the fit.

### Dual luciferase assays

HeLa cells were cultured in DMEM with 10% FBS in 6 well plates seeded at 140,000 cells/ml and incubated at 37oC.

The 5’ UTR sequences (Supplementary Table 3), extending up to the primary ATG initiation codon and including two additional nucleotides (cg) at their 3’ side to maintain the reading frame, were custom synthesized by Azenta Biosciences, Germany. These UTRs were then excised and ligated into the NheI site located upstream of the Renilla luciferase gene in the dual-luciferase vector psiCHECK2 (Promega).

For dual-luciferase reporter assays, HeLa cells were seeded on 6-well plates at a density of 140,000 cells per well. After overnight incubation, reaching ∼60% confluency, cells were transfected with 500ng of the luciferase reporter constructs using the Qiagen Effectene kit, following the manufacturer’s instructions. Twenty four hours post-transfection, cells were washed with 1X PBS and lysed with 1X Passive Lysis Buffer (Promega). *Renilla* and *Firefly* luciferase activities were measured using the Dual-Glo Luciferase Assay System (Promega), according to the manufacturer’s instructions. *Renilla* luciferase activity was normalised to *Firefly* luciferase activity. Samples were analysed with the Promega GloMax Turner 20/20n Luminometer and the data analysed using R. Statistical significance was assessed using ANOVA and Tukey multiple comparison tests.

RNA extractions from cells were performed using TRI reagent (invitrogen) as per the manufacturer’s phase-separation protocol. One microgram of total RNA was reverse transcribed using oligo dT and M-MLV reverse transcriptase (Invitrogen), according to manufacturer’s instructions. No-RT controls were included in parallel. qPCR was performed using the BioRad CFX Connect Real-Time System. cDNA was diluted at a 1:10 ratio, and 5 µl of the diluted cDNA were used in a total reaction volume of 14 µl. Relative mRNA expression was calculated using the 2^(−ΔΔCt) method. Primers used for RT-qPCR are listed in Supplementary Table 4.

## Acknowledgements

This research was made possible through access to data in the National Genomic Research Library, which is managed by Genomics England Limited (a wholly owned company of the Department of Health and Social Care). The National Genomic Research Library holds data provided by patients and collected by the NHS as part of their care and data collected as part of their participation in research. The National Genomic Research Library is funded by the National Institute for Health Research and NHS England. The Wellcome Trust, Cancer Research UK and the Medical Research Council have also funded research infrastructure

We gratefully acknowledge the participants of the National Genomic Research Library (NGRL), whose contributions made this research possible. Secure access to the NGRL under project ID 354 was provided by Genomics England, which delivers the NGRL in partnership with NHS England, and is wholly owned by the UK Department of Health and Social Care. The NGRL contains participants’ health data collected by the NHS as part of their care, along with samples and data from their participation in research, for which fully informed consent has been obtained. This includes genomic and clinical data provided through the NHS Genomic Medicine Service, as well as data obtained through research studies, including the 100,000 Genomes Project and the Generation Study, both of which are delivered in partnership with the NHS, and from other research cohorts involving external collaborators.

With heartfelt thanks to Izzy Barraclough and Dani Breitinger-Blatt for their invaluable support.

---

**Supplementary Figure 1:**
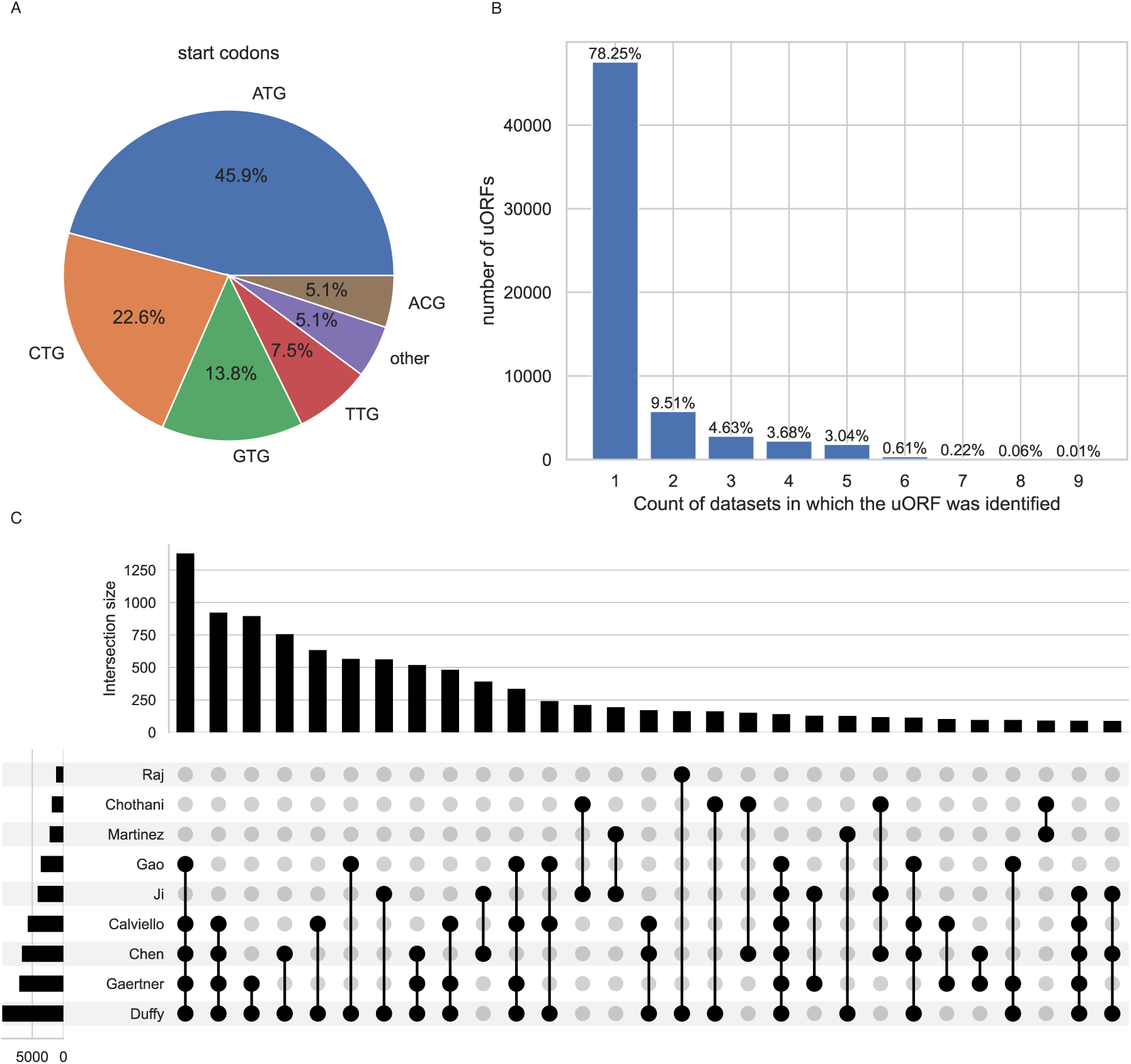
Overview of the uORF data collated in this pipeline. **A)** Proportion of codon sequences identified as the start site of a uORF. **B)** The number of sources which identify each uORF. **C)** Intersection between the sources (minimum subset of uORFs = 80).

**Supplementary Figure 2:**
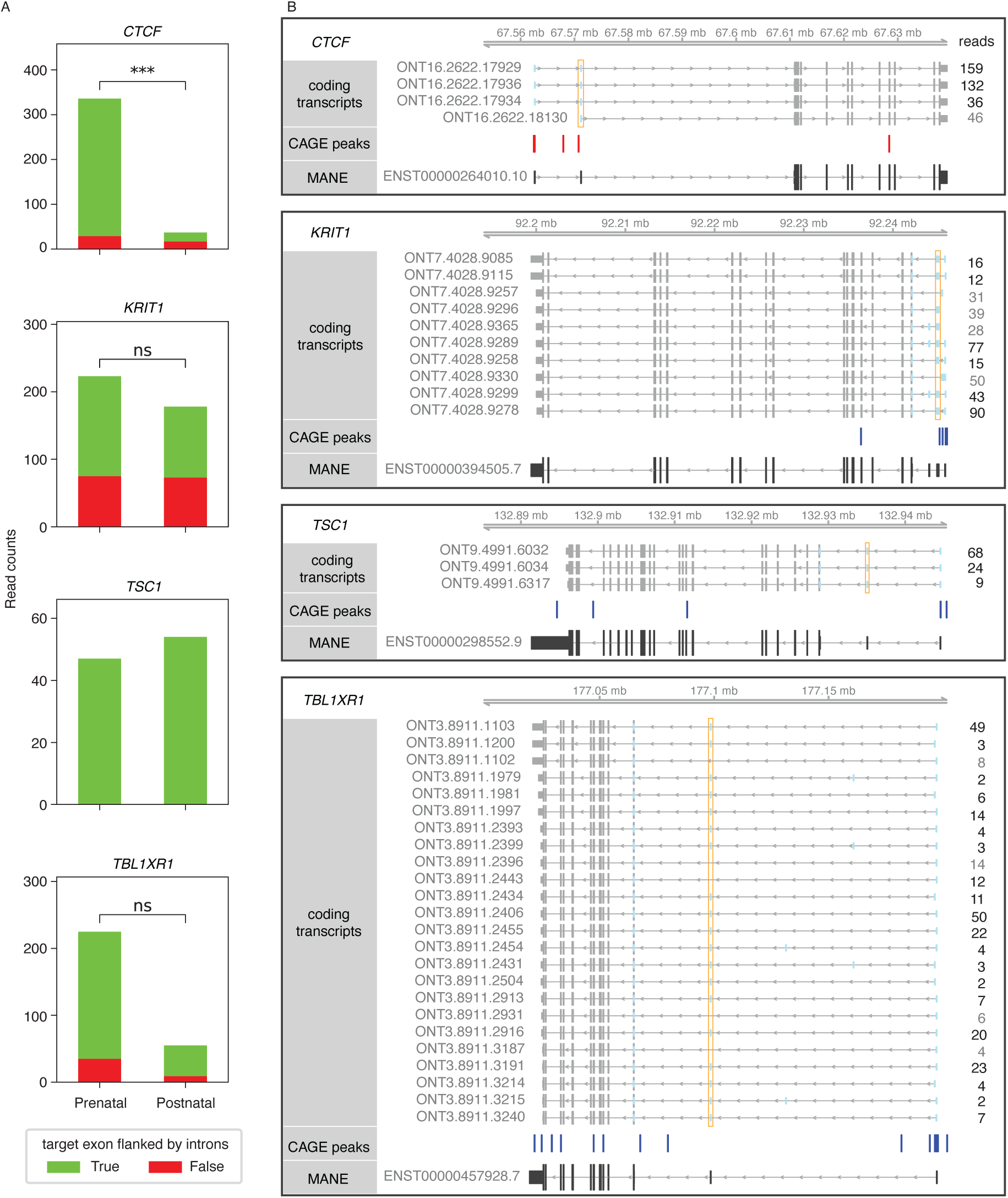
For selected genes **A)** the number of reads from transcripts categorised by whether the target exon is skippable, flanked by introns, in prenatal and postnatal samples. **B)** Isoforms identified by long-read sequencing, predicted to be coding^27^, and supported with a CAGE peak. For *CTCF* only there was a significant decrease in the proportion of reads from transcripts where the target exon can be skipped between prenatal and postnatal samples (fisher exact test, odds ratio = 0.11, p < 0.001).

**Supplementary Figure 3:**
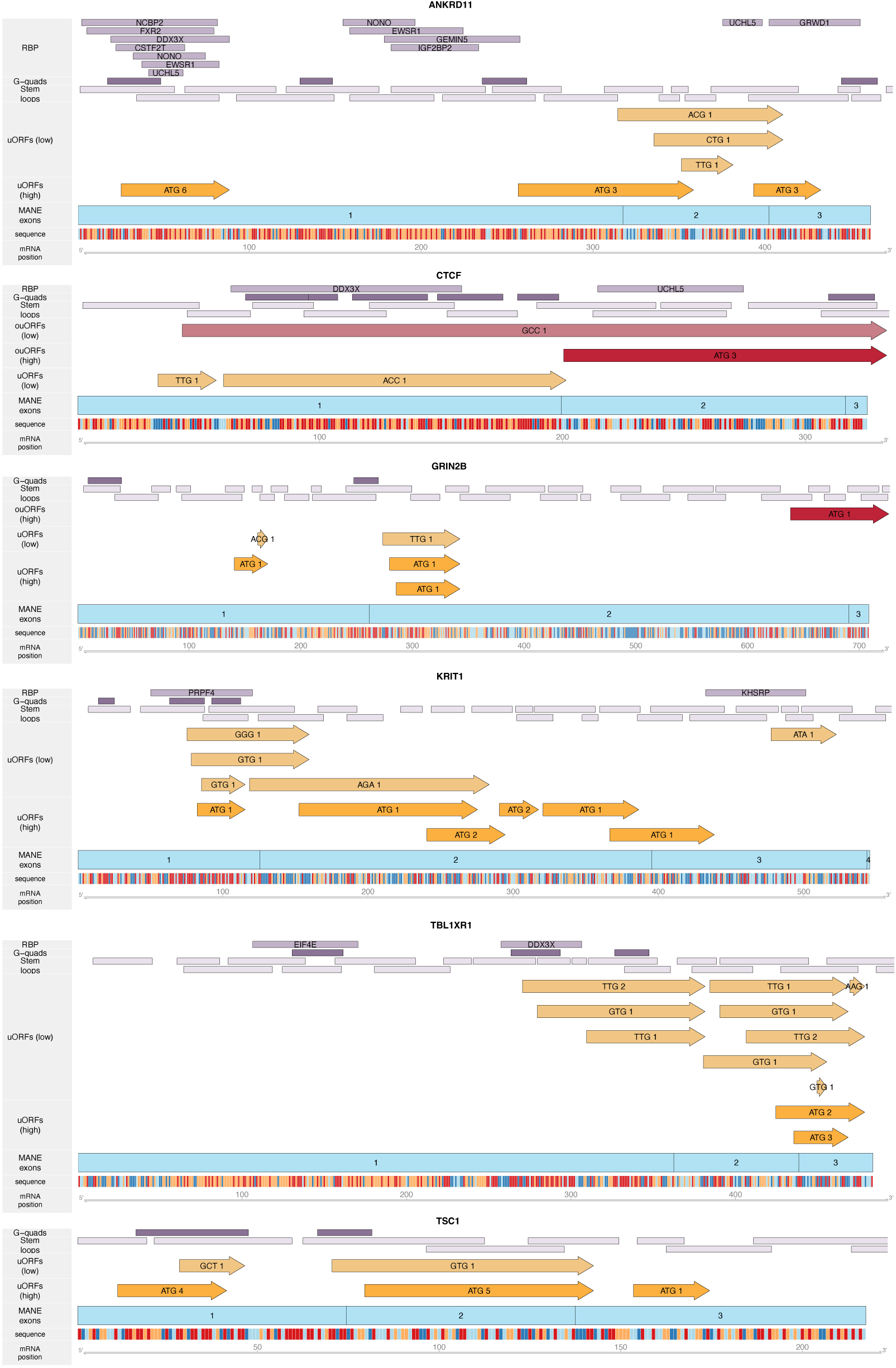
Schematics of 5’UTR features in prioritised gene targets. 5UTR exons are light blue, uORFs are annotated in orange, ouORFs are annotated in red, start codons are highlighted along the sequence. G-quadruplex sections, stem loops and RBPs are annotated in purple-grey. There are no known IRES sites in these six target genes.

**Supplementary Figure 4:**
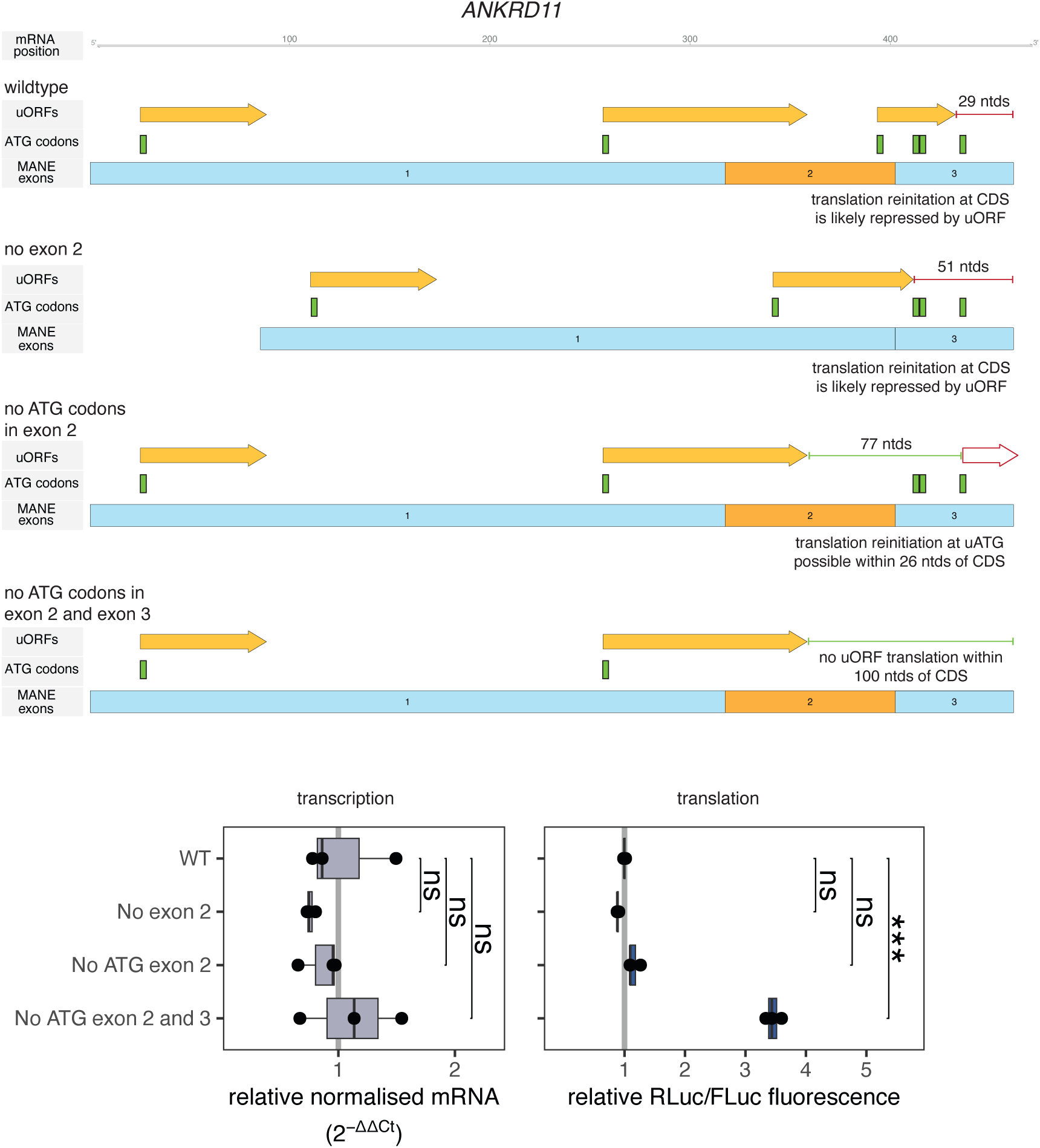
Redundancy between uORFs and normally untranslated uATGs in *ANKRD11* and the impact on exon skipping. In *ANKRD11* the uORF initiating in the target exon has an intercistronic region of 29 ntds which is unlikely to facilitate translation reinitiation at the CDS. However, when the target exon is skipped the uORF initiating in exon 1 extends, such that there is only a marginal increase in the intercistronic region. Further, when the ATG site is mutated, reinitiation may occur at ATGs within exon 3. It is only when the uORF in exon 2, and the ATG codons downstream are removed that translation from the CDS increases.

